# Characterising the methylome of *Legionella longbeachae* serogroup 1 clinical isolates and assessing geo-temporal genetic diversity

**DOI:** 10.1101/2020.09.11.292755

**Authors:** S Slow, T Anderson, DR Murdoch, S Bloomfield, D Winter, PJ Biggs

## Abstract

*Legionella longbeachae* is an environmental bacterium that is commonly found in soil and composted plant material. In New Zealand (NZ) it is the most clinically significant *Legionella* species causing around two-thirds of all notified cases of Legionnaires’ disease. Here we report the sequencing and analysis of the geo-temporal genetic diversity of 54 *L. longbeachae* serogroup 1 (sg1) clinical isolates that were derived from cases from around NZ over a 22-year period, including one complete genome and its associated methylome.

Our complete genome consisted of a 4.1 Mb chromosome and a 108 kb plasmid. The genome was highly methylated with two known epigenetic modifications, m^4^C and m^6^A, occurring in particular sequence motifs within the genome. Phylogenetic analysis demonstrated the 54 sg1 isolates belonged to two main clades that last shared a common ancestor between 108 BCE and 1608 CE. These isolates also showed diversity at the genome-structural level, with large-scale arrangements occurring in some regions of the chromosome and evidence of extensive chromosomal and plasmid recombination. This includes the presence of plasmids derived from recombination and horizontal gene transfer between various *Legionella* species, indicating there has been both intra-species and inter-species gene flow. However, because similar plasmids were found among isolates within each clade, plasmid recombination events may pre-empt the emergence of new *L. longbeachae* strains.

Our high-quality reference genome and extensive genetic diversity data will serve as a platform for future work linking genetic, epigenetic and functional diversity in this globally important emerging environmental pathogen.

**Author Summary:** Legionnaires’ disease is a serious, sometimes fatal pneumonia caused by bacteria of the genus *Legionella*. In New Zealand, the species that causes the majority of disease is *Legionella longbeachae*. Although the analyses of pathogenic bacterial genomes is an important tool for unravelling evolutionary relationships and identifying genes and pathways that are associated with their disease-causing ability, until recently genomic data for *L. longbeachae* has been sparse. Here, we conducted a large-scale genomic analysis of 54 *L. longbeachae* isolates that had been obtained from people hospitalised with Legionnaires’ disease between 1993 and 2015 from 8 regions around New Zealand. Based on our genome analysis the isolates could be divided into two main groups that persisted over time and last shared a common ancestor up to 1700 years ago. Analysis of the bacterial chromosome revealed areas of high modification through the addition of methyl groups and these were associated with particular DNA sequence motifs. We also found there have been large-scale rearrangements in some regions of the chromosome, producing variability between the different *L. longbeacahe* strains, as well as evidence of gene-flow between the various *Legionella* species via the exchange of plasmid DNA.

## Introduction

In both natural and man-made environments *Legionella spp*. bacteria are ubiquitous intracellular parasites of protozoa. Humans are “accidental hosts” when lung macrophages become infected following exposure to contaminated materials, causing Legionnaires’ disease (LD), an often severe form of pneumonia. In New Zealand (NZ), which has the highest reported incidence of LD in the world [1, 2], *Legionella longbeachae* is the species responsible for causing nearly two-thirds of all notified cases [2-4]. Of the two serogroups, serogroup 1 (sg1) is the most clinically relevant. Unlike *L. pneumophila*, the predominant disease causing species in the UK, Europe and USA [1, 5], *L. longbeachae* is primarily found in soil and composted plant material [6, 7]. As a result, most cases of LD in NZ occur over the spring and summer seasons when people at greatest risk are those involved in gardening activities, particularly following exposure to potting mixes and compost [2, 4, 7-9].

Although, whole genome sequencing and interrogation of the resulting data provides invaluable insights into the biology, evolution and virulence of pathogenic organisms, until recently, genomic data for *L. longbeachae* has been sparse, consisting of a single complete genome from an Australian sg1 isolate (NSW150) and four draft genomes; two sg1 and two sg2 isolates [10, 11]. As a result, sequence and comparative genomic analyses between *L. longbeachae* and other *Legionella* species has been limited, with many using the complete NSW150 genome as the sole *L. longbeachae* representative [12, 13]. Despite this, such analyses revealed a larger (≈ 500 kb) and differently organised chromosome than *L. pneumophila* with numerous genes that contribute to its virulence and reflect its soil habitat [10-12, 14].

The recent emergence of *L. longbeachae* as an important cause of LD in Europe and the UK [15], particularly a 2013 outbreak in Scotland [16], prompted concerted efforts in genomic sequencing. This has resulted in a substantial increase in the amount of available genomic data with a large-scale sequencing project of 64 clinical and environmental isolates being reported in 2017 [17]. The availability of a much larger amount of sequence data has revealed further complexity in the *L. longbeachae* genome; variation is driven largely by extensive intra-species horizontal gene transfer and recombination, and there is evidence that there is inter-species gene transfer via plasmids that are the result of recombination between various plasmids of the different *Legionella* species. [17]. Two more complete genomes have also been sequenced, including for the ATCC type strain from one of the first reported cases of LD caused by *L. longbeachae* in Long Beach, California in 1980 (GenBank: FDAARGOS_201; [18]) and one we have obtained from a NZ patient hospitalised with LD in 2014 (F1157CHC; GenBank NZ_CP020894; [19]) that was sequenced as part of this study.

Although the available genomic data to date have provided valuable information about the biology [10] and genetic diversity [17] of *L. longbeachae*, there is scope for further large-scale genomic sequencing in order to more fully assess genetic relationships, determine changes over time and define regions that are important for virulence and pathogenesis. Given the clinical significance of *L. longbeachae* in NZ, particularly the sg1 strain, we have obtained the genome sequence of 54 sg1 clinical isolates, including one complete NZ reference genome [19] and its associated epigenomic data. The isolates were derived from non-outbreak LD cases from 8 regions around NZ over a 22-year period (1993-2015), allowing us to examine its geo-temporal genetic diversity through ancestral state reconstruction and phylogenic analysis, and for the first time, characterise its epigenome.

## Results & discussion

### Genome architecture

#### Chromosome

Given there are few complete *L. longbeachae* genomes available we chose one of our sg1 isolates as the reference genome to further analyse our other 53 NZ isolates. This isolate, called F1157CHC, was sequenced using both Illumina short read sequencing in the initial comparative dataset, and then subsequently sequenced with PacBio long read sequencing. The complete chromosome has been recently published [19], and therefore the description of this genome is kept relatively brief and is more comparative in nature with the other available reference *L. longbeachae* genomes (NSW150, Genbank accession no. NC_013861; FDAARGOS_201, Genbank accession no. NZ_CP020412).

We compared all three reference genomes using the MAUVE plugin within Geneious (v 9.1.8) and the results of this analysis are shown in Fig 1. At 4,142,881 bp, F1157CHC is larger by 65,549bp when compared to NSW150 and smaller by 19,851bp to FDAARGOS_201. Overall the genomes of F1157CHC, NSW150 and FDAARGOS_201 are very similar in their organisation with the MAUVE alignment showing four (81 – 2,264kb) collinear blocks in the genomes, hereafter called LCB1, LCB2, LCB3 and LCB4 by visualisation in Geneious, but does not reflect their genomic order. At an overall genome level, the order and orientation of these blocks indicates a greater similarity between NSW150 and F1157CHC, while FDAARGOS_201 is slightly different (S1 Table).

**Fig 1:**
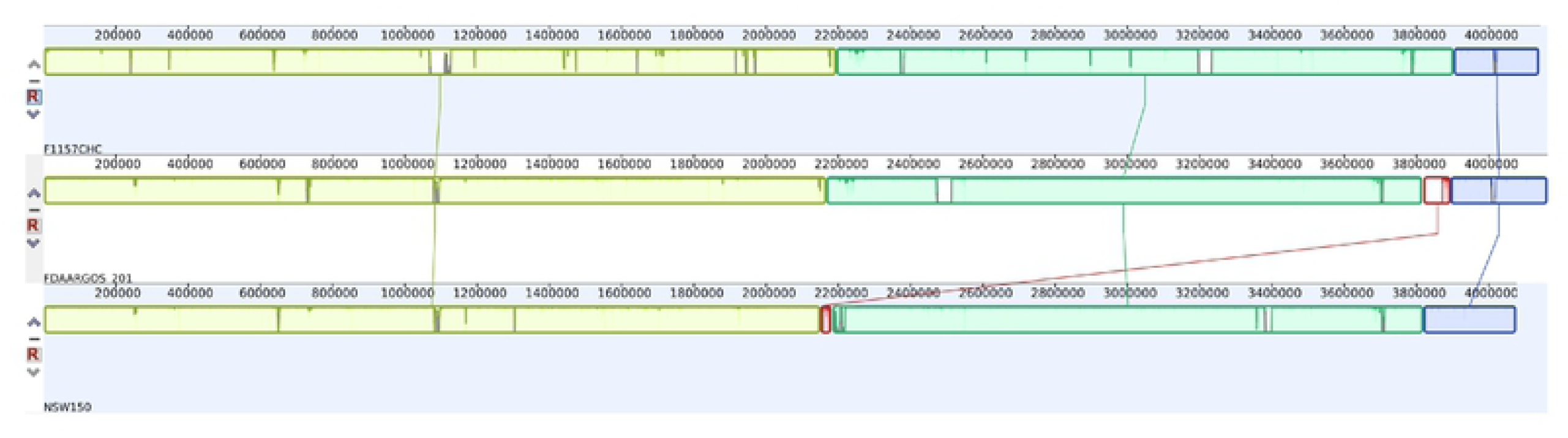
Mauve alignment of the three complete *L. longbeachae* sg1 genomes, F1157CHC, FDAARGOS and NSW150 from top to bottom. The 4 main collinear blocks are indicated by colours (LCB1 is red, LCB2 is yellow, LCB3 is green and LCB4 is blue). The sizes of the blocks for LCB1, LCB2, LCB3 and LCB4 are ∼81kb, ∼2265kb, ∼1807kb and ∼272kb respectively. Lines between LCBs in the isolates are for the same LCB. In the visualisation the areas within the LCBs without colour indicates underlying differences in the LCB, as explained by the fact that the LCBs are different sizes in each isolate.

Three of these blocks (LCB2, LCB3 and LCB4) are found in all three genomes, and a further one (LCB1) is found only in NSW150 and FDAARGOS_201. The genomic coordinates and the percentage of the collinear block that contains genomic sequence are described in S1 Table. In addition, there are two and three small regions in two of the genomes that are not found in collinear blocks totaling 4.2 and 4.4kb for NSW150 and FDAARGOS_201, respectively. For FDAARGOS_201 and NSW150, two of these unique regions are found flanking the shortest collinear block of 81kb (LCB1), and for NSW150 the third region is a short sequence at the start of the chromosome (unusually for this chromosome the start of the *dnaA* gene is not annotated to be at position 1 of the chromosome). The LCB1 block shows the greatest disparity in content with the genomic length in NSW150 being 31.3kb, whereas it is 73.6kb in FDAARGOS_201, hence there are many gaps that are incorporated into the collinear block alignments.

It should be noted that as the MAUVE aligner within Geneious works on a linear chromosome, the LCBs at the end of the chromosome form part of the same larger collinear block, meaning that on the circular chromosomes for these three *L. longbeachae* isolates there are in effect only three blocks, with the block of ∼1807kb (LCB3) being flanked by the content variable 81kb block LCB1. There are thus only a few boundaries around the main collinear blocks. The boundary between LCB2 and LCB3 in FDAARGOS_201 and FH1157CHC occurs within the traF gene, part of the *tra* operon. The organization is more complex in NSW150 in that the 31.5kb block of LCB1 and a 3.9kb region containing three hypothetical genes is found between LCB2 and LCB3, with the *tra* operon being found on LCB1. The *tra* operon forms part of the Gram-positive type IV secretion system (T4SS) for the transfer of plasmids via conjugation [20], so it is therefore an important operon for pathogenicity. For the other main boundary between LCB3 and LCB4, the transfer messenger RNA (tmRNA) *ssrA* gene is present at the end of LCB4 for all three chromosomes. The tmRNA genes are part of the trans-translocation machinery which can overcome ribosome stalling during protein biosynthesis. Trans-translocation has been found to be essential for viability within the *Legionella* genus, with the *ssrA* gene being unable to be deleted in *L. pneumophila* [21]. Under the control of an inducible promoter, it was found that decreasing tmRNA levels led to significantly higher sensitivity to ribosome-targeting antibiotics, including erythromycin [21]. At the end of LCB3 in FH1157CHC and NSW150, there is an IS6 family transposase and an SOS response-associated peptidase, about which little is known for either of these genes. The flanking gene in FDAARGOS_201 comes from a small orphan block of 1.3kb between LCB1 and is a short DUF3892 domain-containing protein, as defined by Pfam [22]. Whilst being of unknown function this gene is found widely across bacteria and archaea, and within the Legionellales order.

As described above, the collinear blocks defined by MAUVE include gaps, and except for LCB1, all other defined blocks in the isolates are found with the genomic length being greater than 87% of the block length. Within the blocks themselves, LCB1 shares a common region of ∼23.8kb and a larger non-overlapping (i.e. different gene content) region in FDAARGOS_201 compared to NSW150. For the remaining three blocks there are combinations of absence and presence of genetic material within these blocks and across the three isolates. For the regions over 10kb, these can be summarized as regions that are present in only a single isolate (37.1kb in LCB3 of F1157CHC), or in two isolates (12.0kb in LCB2 of FDAARGOS_201 and NSW150), and those that are different across all three isolates (86.3kb in LCB2, 12.2kb in LCB2 and 18.1kb in LCB3). Other more complex combinations of the three isolates account for a further six regions of the blocks involving three regions for F1157CHC and NSW150 (19.3kb in LCB2 with a differing FDAARGOS_201 sequence; 64.4kb in LCB3 with an extra sequence in NSW150; and 43.4kb in LCB3, a large FDAARGOS_201 sequence), two for FDAARGOS_201 & NSW150 (40.0kb in LCB4 with different flanking sequence for FDAARGOS_201 and F1157CHC; and 11.3kb in LCB2 with a different sequence for F1157CHC), and one for FDAARGOS_201 & F1157CHC (101.3kb in LCB3 with a different sequence for NSW150). The gene content in these blocks is varied, and the boundaries can be close to tRNA genes, site-specific integrase genes, SDR family oxidoreductase genes, ankyrin repeat domain-containing genes, or in intragenic space, but for some of the boundaries transposase genes (IS3, IS4, IS6, and IS926 families) are involved. In bacteria, tRNAs have been shown to be integration sites [23], so finding them at boundaries of the defined collinear blocks may not be all that surprising, especially given the presence of integrases at these locations.

In order to visualise the data from the comparison of the 54 samples in the dataset and to show multiple facets of this study simultaneously, an overarching Circos figure (Fig 2) was generated using the complete PacBio genome of F1157CHC as a backbone. The tracks in the figure are described in detail in the figure legend. Overall it can be seen that the regions detected for recombination by Gubbins are unevenly distributed around the genome, with some clusters around the genome (∼600kb, ∼800kb, ∼1900 – 2050kb), and large, slightly less dense region (2300 – 2800kb).

**Fig 2:**
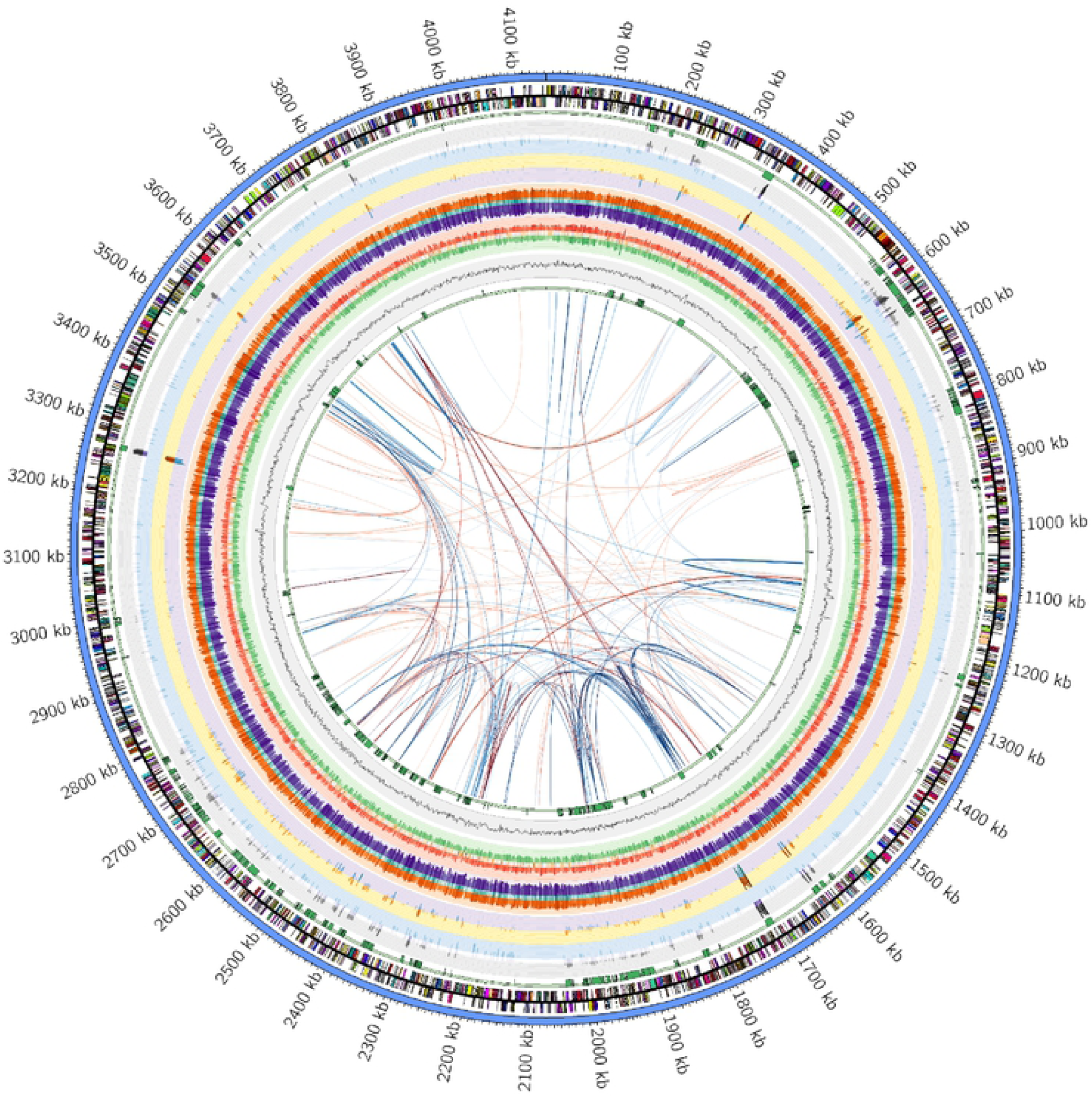
Circos plot of NZ *L. longbeachae* isolate F1157CHC. Tracks from the outside to the inside are; chromosomal ideogram (blue), genes on the plus strand, genes on the minus strand (both annotated via eggNOG), regions of recombination detected by Gubbins in the full NZ dataset of 54 sg1 isolates, a histogram showing Snippy-detected SNPs in recombination areas, a heatmap of all SNPs, a histogram showing Snippy-detected SNPs in non-recombination areas, a histogram of non-synonymous SNPs, a histogram of synonymous SNPs, a histogram of m4C densities consistent with gene strand, a heatmap of all m4C densities, a histogram of m4C densities inconsistent with gene strand, a histogram of m6A densities consistent with gene strand, a heatmap of all m6A densities, a histogram of m6A densities inconsistent with gene strand, a line plot of the GC percentage and finally a repeat of the regions of recombination detected by Gubbins in the full NZ dataset of 54 sg1 isolates. For the gene predictions, the genes are coloured by functional COG category and the colours used are described in Supplementary Figure 1. All data for the SNPs, methylation patterns and GC percentage are values calculated in non-overlapping 1kb bins. The data for the SNPs and methylation patterns are shown in a log10 scale, and the histograms are also coloured so that larger values are in darker colours. In the centre of the plot are the results from the Reputer analysis, with palindromic repeats in red and forward repeats in blue. The repeats are darker in colour with a smaller Hamming distance between the repeats.

#### Plasmid

Of the three available reference genomes, only NSW150 and our genome F1157CHC were found to contain a plasmid (pNSW150, Genbank accession no. NC_014544; pLLO_F1157CHC, Genbank accession no. CP020895; [19]). At 108,267 bp pLLO_F1157CHC is 36,441 bp larger when compared to pNSW150. To assess plasmid architecture more fully, we identified three additional *L. longbeachae* plasmids through further sequencing of two of our clinical isolates, B1445CHC and B41211CHC. Isolate B1445CHC was found to contain two plasmids of 73,728 bp and 150,426 bp (pB1445CHC_73k; Genbank accession CP045304; and B1445CHC p150k; GenBank accession CP045305), while B41211CHC contained only one plasmid of 76,153 bp (pB41211CHC_76k; GenBank accession CP045307). All five *L. longbeachae* reference plasmids were aligned using MAUVE and visualised in Geneious, with the results shown in Fig 3. The plasmids share a common backbone consisting of conjugational genes (yellow collinear block), which range in size from ∼25,000 to ∼28,000 bp, as well as several other collinear blocks that vary in size and orientation (Fig 3). These blocks are separated by variable regions around mobile genetic elements, such as insertion sequences. Similarly, analysis of the larger plasmid pB1445CHC_150k revealed this is the same as plasmid pLELO (Genbank accession NC_018141), which was first reported in *L. pneumophila*. MAUVE alignment of the *L. longbeachae* plasmids, pLELO and two *L. sainthelensi* plasmids (LST pLA01-117_165k Genbank accession CP025492; LST pLA01-117_122k Genbank accession CP025493, [24]) (S2 Fig) again shows the *Legionella* plasmids have a common backbone including conjugational genes (yellow collinear blocks) separated by variable regions. Although the number of plasmids in our analysis is limited, the data also show that the *Legionella* plasmids identified to date can be broadly divided into two groups; one that consists of the smaller plasmids of around ∼70kb that appear to be primarily a *L. longbeachae* group (pNSW150, pB1445CHC_73k, pB41211CHC_76k) and another group consisting of the larger plasmids that occur in various species, including our *L. longbeachae* complete genome (pLLO_F1157CHC, pLELO LST, pLA01-117_165k). Combined these data suggest there has been extensive plasmid recombination as well as intra-species and inter-species transfer, and plasmids may be an important means of exchanging genetic material both between and within different *Legionella* species, supporting the findings of Bacigalupe et al., [17].

**Fig 3:**
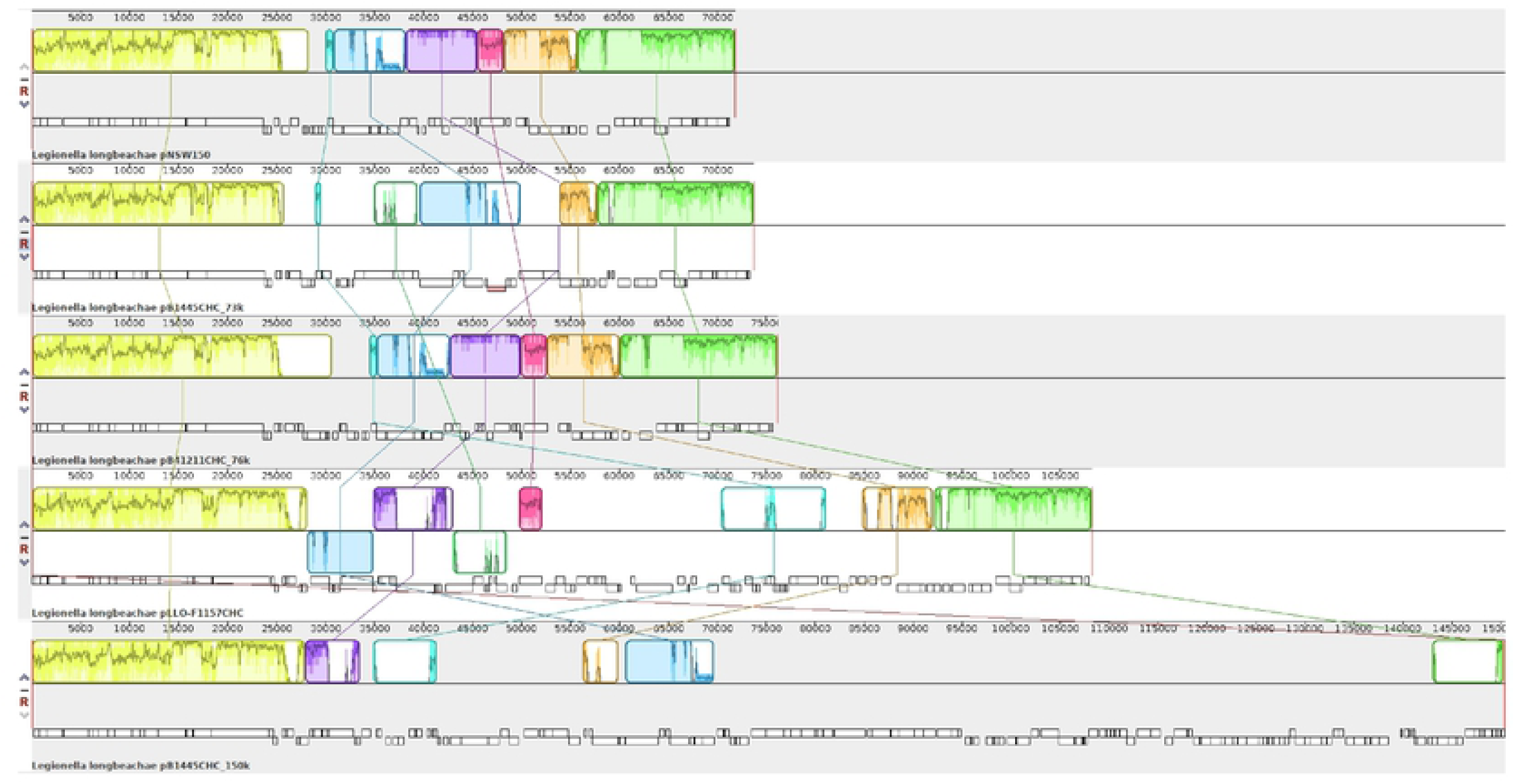
Mauve alignment of five *L. longbeachae* reference plasmids

Interestingly, pB1445CHC_73 k has a repetitive region that has identified as a clustered regularly interspaced short palindromic repeat (CRISPR) element. This element belongs to the type I-F system with the same repeat region between 20 to 33 spacer regions and cas1-cas6f associated enzyme (Fig 4). While there are few reports of naturally occurring CRISPR-Cas arrays on plasmids, previous studies [25, 26] as well as a recent comparative genomics analysis of all publically available bacterial and archaeal genomes has demonstrated that type IV CRISPR-Cas systems are primarily encoded by plasmids [27]. To our knowledge, our findings are the first report of a type I-F CRISPR-Cas array being present on a plasmid. Further analysis of the other *L. longbeachae* isolates sequenced as part of this study showed that this CRISPR element is also found in 5 other strains reported here (F2519CHC, LA01-195, LA01-196, LA03-576).

**Fig 4:**
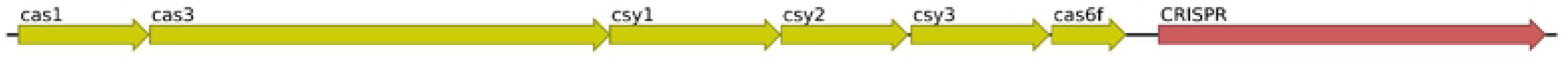
The type I-F CRISPR-Cas element found in some of the *L. longbeachae* isolates sequenced in this study

### Genetic diversity of *L. longbeachae* sg1 clinical isolates

#### Gene content and functional gene categories

As indicated in the gene rings in Fig 2, the genome of F1157CHC was functionally annotated and categorized using the amino acid sequences from the NCBI PGAP predictions against the eggNOG-mapper database (v. 2.0). The genes were then coloured according to their COG categories (S1 Fig). We found that for the 3,622 predicted genes, 3,410 (94.14%) annotations resulted, and of those 2,741 (80.38%) were categorized with COG functional categories meaning that 669 (19.61%) genes were not annotated. In terms of the performance of the eggNOG server, this level of annotation for *L. longbeachae* is slightly above the level of 76% reported for the Legionellales order, and close to the eggNOG v. 5.0 database average of 80% [28]. The main functional groups i.e. those with a single COG category definition account for 2,522 (73.96%) of the annotations. COG category S – “function unknown” – is the largest single category, and accounts for 547 (16.04%) of the returned annotations, though with the output, all genes for which a COG category has been defined have a gene description. The eggNOG output can be found in S2 Table.

We used our set of 53 draft genomes to investigate both the core and pangenome of the isolates in the BioProject PRJNA417721. A genome summary of these 53 draft genomes, plus the complete PacBio F1157CHC and the NSW150 and FDAARGOS_201 complete genomes, can be found in S3 Table. The 53 draft genomes all have a similar length (4148006.3 bp ± 78321.3), GC content (37.11% ± 0.05), coding sequences (3576.2 ± 81.8), signal peptides (277.6 ± 7.8) and tRNAs (45.1 ± 3.0) amongst themselves. These values are all lower compared to the values from the complete genomes (genome length: 4194301.3 bp ± 66793.1; coding sequences: 3609.7 ± 76.8; signal peptides: 282.7 ± 5.5; and tRNAs: 48.3 ± 3.2), except for the GC content which is slightly higher (37.15% ± 0.10). Due to the draft nature of the assemblies, on a proportional basis the biggest difference is seen in the number of rRNAs (6.89 ± 0.42) which is much less than the 12 found in the complete genomes, but this is to be expected with repetitive DNA sequences and short read sequencing.

In comparison to the recent study of Bacigalupe et al., [17] covering sg1 isolates from mostly Scotland (n = 50), as well as the USA (n = 1), Australia (n = 1) and New Zealand (n = 4), we found that the range of our coding sequences was not different to the 3,558 genes they reported. However, as only summary gene numbers per genome were provided, we cannot say if there is any real difference in gene numbers between the NZ strains reported here, and the primarily Scottish strains in the Bacigalupe et al. study [17], but it seems unlikely to be the case.

We used Roary [29] to analyse the core and pangenome of the 56 *L. longbeachae* isolates. We found a pangenome of 6,517 genes, and a core genome of 2,952 genes, which in this case indicates the number of genes present in all 56 isolates, including the reference genomes. This number is therefore ∼86.3% of the number of genes in the F1157CHC genome, indicating a large core genome and a small accessory genome amongst the *L. longbeachae* isolates in this study. Given the geotemporal collection of our study, within a 22-year period in New Zealand that may not be a surprising finding that an average of 468 genes were found in the accessory genome. Bacigalupe et al., [17] also reported a core genome (2,574 genes) and pangenome (6,890 genes) for the 56 *L. longbeachae* sg1 strains they studied, which were over a shorter, but contemporaneous, timeframe. Given the isolate numbers are almost the same (excluding reference isolates), but that the methodologies for calculating the core genome were different, it is interesting to observe a smaller number of genes in the core but a larger number of genes in the pangenome. It is tempting to speculate that there might be a smaller gene repertoire for *L. longbeachae* in New Zealand, again as a result of its relative geographical isolation, or maybe that environmental conditions are different, requiring the use of different sets of genes to survive within the New Zealand soil. Using the categories defined within Roary, we found 157, 865 and 2,543 genes in the soft core (95 to 99% of strains), shell (15 to 95%) and cloud (0 to 15%) genomes respectively. The Roary output is shown in S1 File.

Currently, there are 61 recognised species and 3 subspecies within the *Legionella* genus (http://www.bacterio.net/legionella.html). Of these, 58 species have had at least draft genome sequencing performed on isolates or type strains in order to understand the evolution of the genus [30], for which a core genome has been estimated to be only 1,008 genes, highlighting the diversity within the genus. *L. pneumophila* is regarded as the most clinically important pathogen within the genus [1] with a GC content of ∼39% and a smaller genome of ∼3.3Mb. A recent Australian study [31] has estimated the core genome of this species to be 2,181 genes, with a pangenome of 5,919 genes, representing a 36.7% fraction of all genes in the *L. pneumophila* pangenome being in the core genome. In comparison, in our study, analogous numbers indicate a fraction of 45.3% for *L. longbeachae*. This suggests that the *L. longbeachae* genome is probably more stable than the *L. pneumophila* genome.

Finally, we used FastGeP [32] to perform an *ad-hoc* whole genome MLST analysis of the 56 isolates using the 3,420 CDSs in the F1157CHC (CP020894) reference genome. We found that 2,756 loci were shared by the 56 genomic sequences of which 1,321 (47.93%) were identical at the allelic level. One-hundred and eight of the shared loci were excluded because of hypothetical gene duplications, and 664 were excluded because of incomplete information, such as missing alleles, truncation or nucleotide ambiguities. After removal of these loci, 2,648 (of which 1,327 were polymorphic) were used to construct the distance/difference matrix. With the different methodological approach, it is not surprising that this value is different to that calculated by Roary. Visualization of the FastGeP matrix in iTOL [33] is shown in S2 File.

#### Antimicrobial resistance and virulence genes

The 54 *L. longbeachae* sg1 isolates all contained a chromosomal class D β-Lactamase gene that is homologous to *bla*_*OXA*_ enzyme family. This 795 bp *bla*_*OXA-like*_ gene, whose phenotypic features are uncharacterized, is also found in *L. oakridgensis* (100% nucleotide match). Twenty-one isolates also have another molecular class D β-Lactamase with 100% nucleotide match to *bla*_*OXA-*29_ that are contained on a plasmid similar to *L. pneumophila* pLELO. The *bla*_*OXA-*29_ gene was first identified in the *Fluoribacter gormanii* type strain ATCC 33297^T^ (Genbank accession number NG_049586.1; [34]). The majority of the known class D β-Lactamases are found on mobile genetic elements and indicate the intra-species transfer of *bla*_*OXA-*29_ on conjugative plasmids amongst the various *Legionella* species such as *L. pneumophila, L. sainthelensi*, and *L. heckeliae*. This *bla*_*OXA-*29_ β-Lactamase is part of a group of structurally related serine enzymes that have high activity against penicillins, reduced activity against cephalosporins and no activity against carbapenems [35].The *bla*_*OXA*_ genes are located downstream of the *bla*I (transcriptional regulator gene) and accounts for the intrinsic resistance of *Legionella* spp. to the penicillins. All isolates also contained a tetracycline destructase gene, *tet56*, which had previously been identified in *L. longbeachae* through comparative gene analysis by Forsberg et al., [36] and was shown to confer tetracycline resistance when expressed. Tet56 belongs to a recently identified family of flavoprotein monoogenoxygenases that inactivate tetracycline antibiotics through covalent modification to the antibiotic scaffold [36, 37]. Previously, the antimicrobial susceptibilities of 16 isolates that were sequenced in the current study had been investigated [38]. For these isolates, the tetracycline MIC_90_ was found to be high, ranging between 16 to 64 mg/mL when the isolates were grown in BYE broth, suggesting that tet56 was expressed in these isolates and was functional, inactivating the tetracycline (S4 Table).

Virulence factors database analysis showed the 54 isolates had between 33 and 36 virulence factor genes (S5 Table). Many of these encoded various components of the type IVB Dot/Icm secretion system, which has been found to be present in all *Legionella* species examined to date and is essential for its virulence [39].

### Geo-temporal and phylogenetic relationships

#### New Zealand *L. longbeachae* isolates

The 54 *L. longbeachae* sg1 isolates were found to share 5,383 core SNPs (2,338 post Gubbins) and phylogenetic modelling estimated that these isolates contained a substitution rate of 3.58 x 10^−8^ – 1.51 × 10^−7^ substitutions site^-1^year^-1^ (95% HPD interval) and shared a date of common ancestor between the years 108 BCE - 1608 CE (95% HPD interval). The isolates had undergone a large amount of recombination, as evidenced by the large number of recombinant SNPs identified via Gubbins (3,045) and were shown to belong to two main clades (Fig 5). The larger of the two clades consisted of isolates from multiple different regions in New Zealand, while the smaller clade consisted of isolates from the Canterbury district only (S3 Fig). However, as the Canterbury district was oversampled in this study, this finding could just be due to random chance.

**Fig 5:**
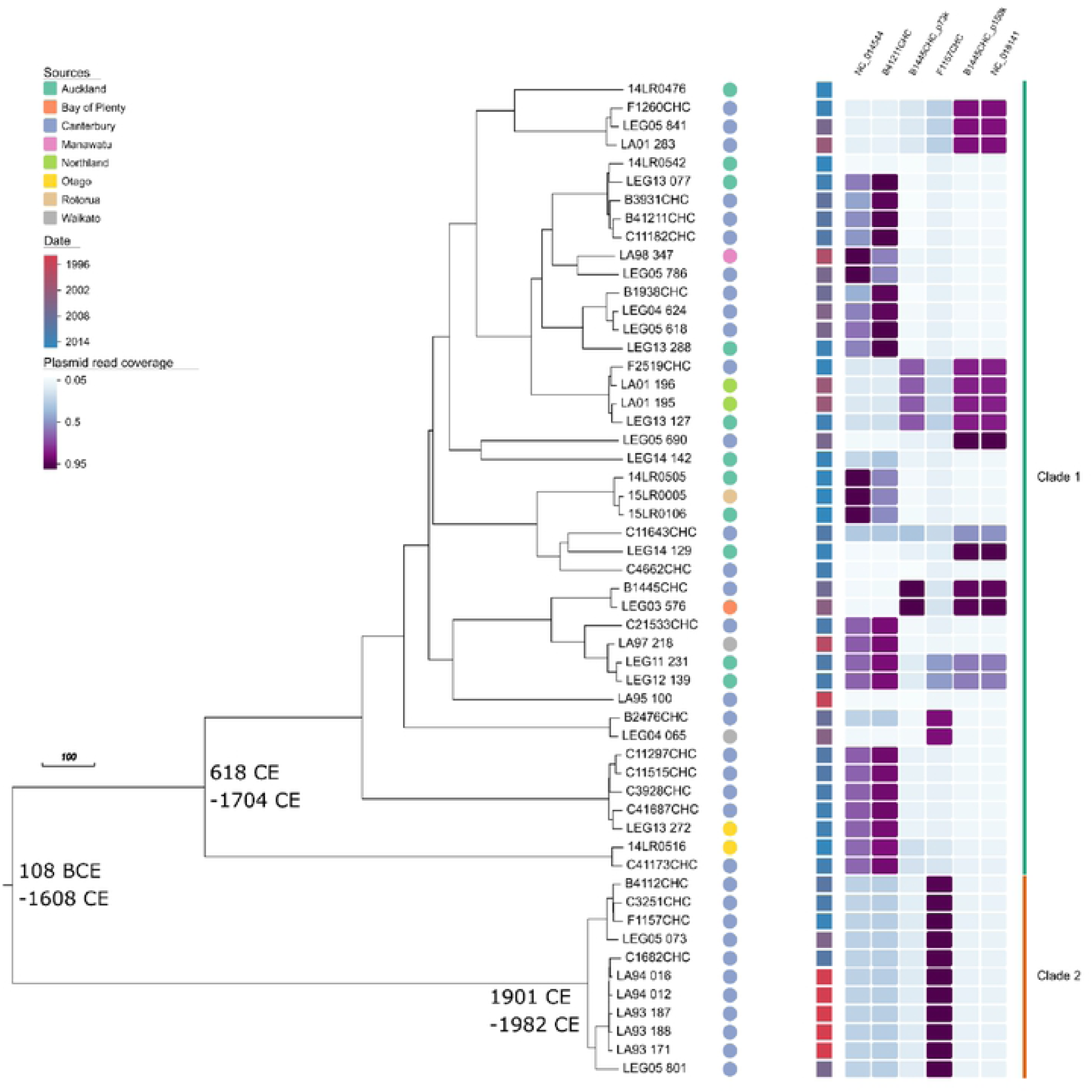
Maximum clade credibility tree of 54 *L. longbeachae* clinical isolates. The scale bar represents the length of 100 years. Isolates are coloured by date of collection (squares), region (circles) and plasmid read coverage (heat map). The years in parentheses represent the estimated timing of coalescent events (95% Highest Posterior Density interval).

Alignment of the New *Zealand L. longbeachae* read sets to the six *Legionella* reference plasmids demonstrated that some of the New Zealand *L. longbeachae* isolates contained an exact copy of the reference plasmids investigated with reads aligning to the entire reference sequence, some contained similar plasmids with reads aligning to sections of the reference plasmids, and some contained reads that aligned to sections of more than one reference plasmid (Fig 5). This again illustrates that the *Legionella* plasmids share a common back bone separated by variable regions and that there is extensive recombination amongst them (S4 Fig). The plasmid results also correlated with the clades identified via phylogenetic analysis, suggesting that plasmid recombination events may pre-empt the emergence of new *L. longbeachae* strains.

#### Global *L. longbeachae* isolates

The 89 *L. longbeachae* isolates from the United Kingdom and New Zealand shared 3,219 core SNPs and belonged to multiple small clades. Most of the clades consisted only of isolates from a single country, whilst a small number consisted of isolates from both countries (S5 Fig). This indicates some recent global transmission of *L. longbeachae*.

### L. longbeachae methylome

Methylome analysis of our NZ reference genome strain identified two classes of modified base, N4-cytosine (m^4^C) and N6-methyladenine (m^6^A). Bases in the chromosomal sequence were more likely to be modified (1.49% of As and 6.4% of Cs being methylated) than those in the plasmid (1% of As and 2.4% of Cs) (Fig 6A). Modifications were evenly distributed within a given molecule, with the exception of a single cluster of m^6^A in the chromosome (Fig 6B). The majority (73.6%) of m^6^A bases occurred in three sequence motifs (ATGNNNNNNRTGG/CCAYNNNNNNCAT, GATC and GGGAG). Two of these motifs (ATGNNNNNNRTGG/CCAYNNNNNNCAT and GATC) are almost always methylated (97-99.5% of occurrences in the genome) while the third (GGGAG) is frequently modified (77.2% of occurrences). By contrast, the m^4^C modifications are not strongly concentrated in motifs. The motif most frequently associated with this modification (CSNNNTB) is only modified in 9.2% of occurrences in the reference genome (about 3 times the background rate for all cytosines).

**Fig 6:**
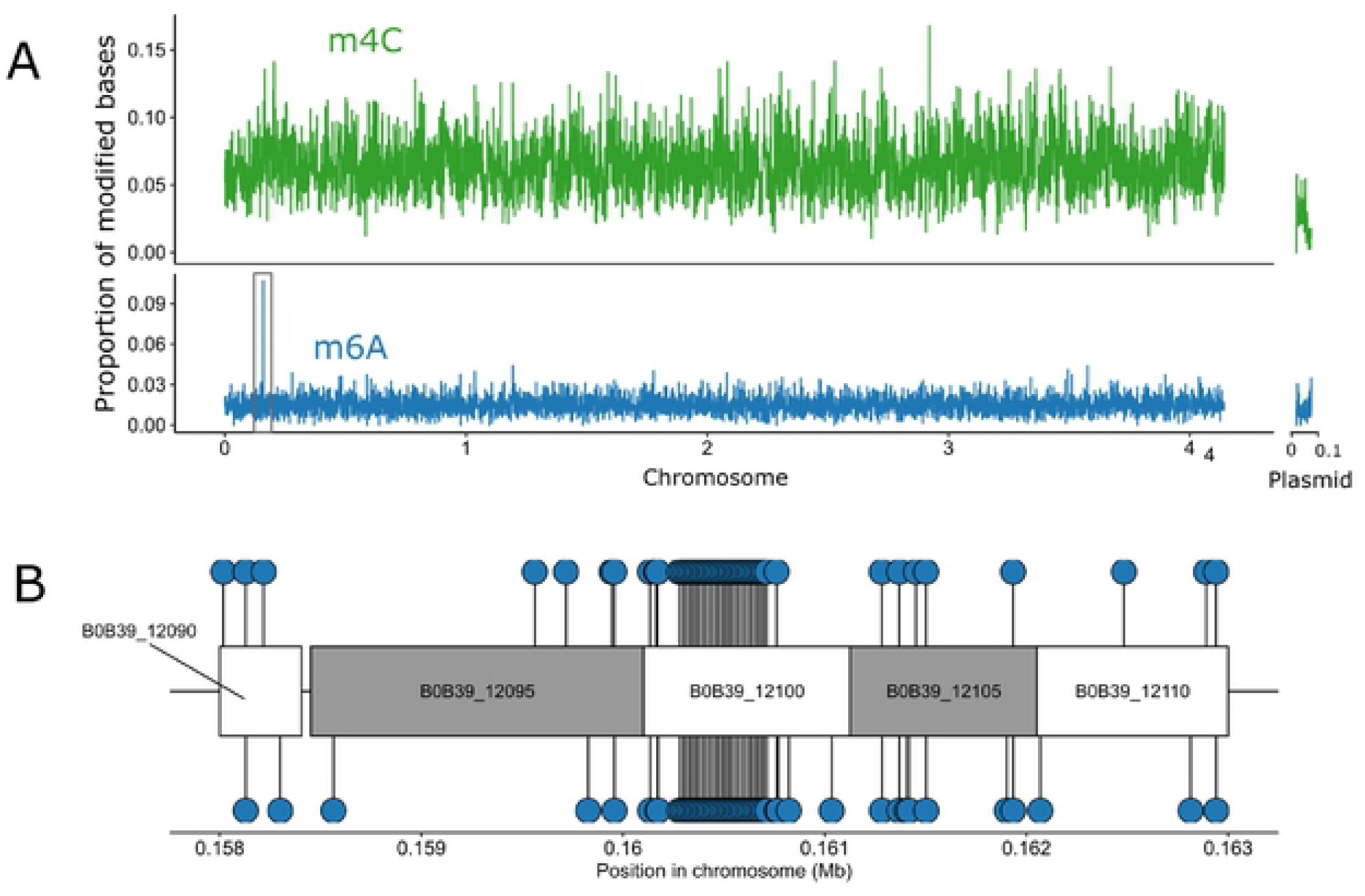
Methyl-distribution. Both methylation marks are approximately evenly distributed across the *L. longbeachae* genome. (A) The frequency with which m4C (above in green) and m6A (below in blue) modifications were detected is plotting in 1kb windows, note the difference in y-axes for each sub-plot. The black box represents the region of unusually high m6A modifcations highlighted in (B). (B) The locations of individual m6A modifications are shown as blue circles for the region with an unsually high rate of this mark. The alternating white and grey boxes represent genes in this region, and are labelled with their NCBI locus ID. The gene responsible for the very high rate of m6A modification, B0B39_12100, is a tetratricopeptide repeat protein.

DNA methylation in bacteria is often associated with restriction modifications (RM) systems, which protect the bacterial cell from foreign DNA. These systems combine a restriction encodnuclease that digests un-methylated copies of a target sequences and a DNA methyltransferase that methylates this sequence motif in the bacterium’s own DNA. The strong association between m^6^A modification and three sequence motifs in the *L. longbeachae* genome suggests this modification is part of an RM system.

Using REBase, we identified putative methyltransferases and encodnucleases in the *L. longbeachae* genome. This analysis revealed three neighbouring genes that encode a type I RM system associated with the ATGNNNNNNRTGG/CCAYNNNNNNCAT motif. Specifically, gene B0B39_08545 encodes a SAM-dependent DNA methyltransferase with target recognition domains for both ends of this motif, while genes B0B39_08550 and B0B39_08555 encode the S and R subunits of an associated endonuclease. The enzymes responsible for the CATC and GGGAG motifs are less clear. Two proteins (LloF1157ORF6795P and LloF1157ORF8795P) are homologous to methyltransferases that recognize GATC in other species. Neither of these proteins are associated with a restriction endonuclease.

The m^4^C modification is not strongly associated with any sequence motif in *L. longbeachae*. Although many bacterial genomes contain this modification, the biological functions encoded by it remain unclear [40]. There is some evidence that this mark may contribute to the regulation of gene expression. Notably, the deletion of a single m^4^C methyltransferase in *Helicobacter pylori* alters the expression of more than 100 genes and leads to reduced virulence. We used our genome annotation and methylation data to tests for any associations between m^4^C methylation and genome features of functional classes of genes that might suggest this mark contribute to gene regulation in *L. longbeachae*. We found this mark is considerably more common within protein coding genes than in intergenic spaces (Fig 7A). However, there is no association between the presence of this mark in a gene sequence and any of the functional classifications present in our COG data (Fig 7B). Although the over-representation of m^4^C bases in genetic sequences suggests this mark might be associated with, or a passive consequence of, transcription in *L. longbeachae*, we find no evidence that this mark contributes to particular biological functions.

**Fig 7:**
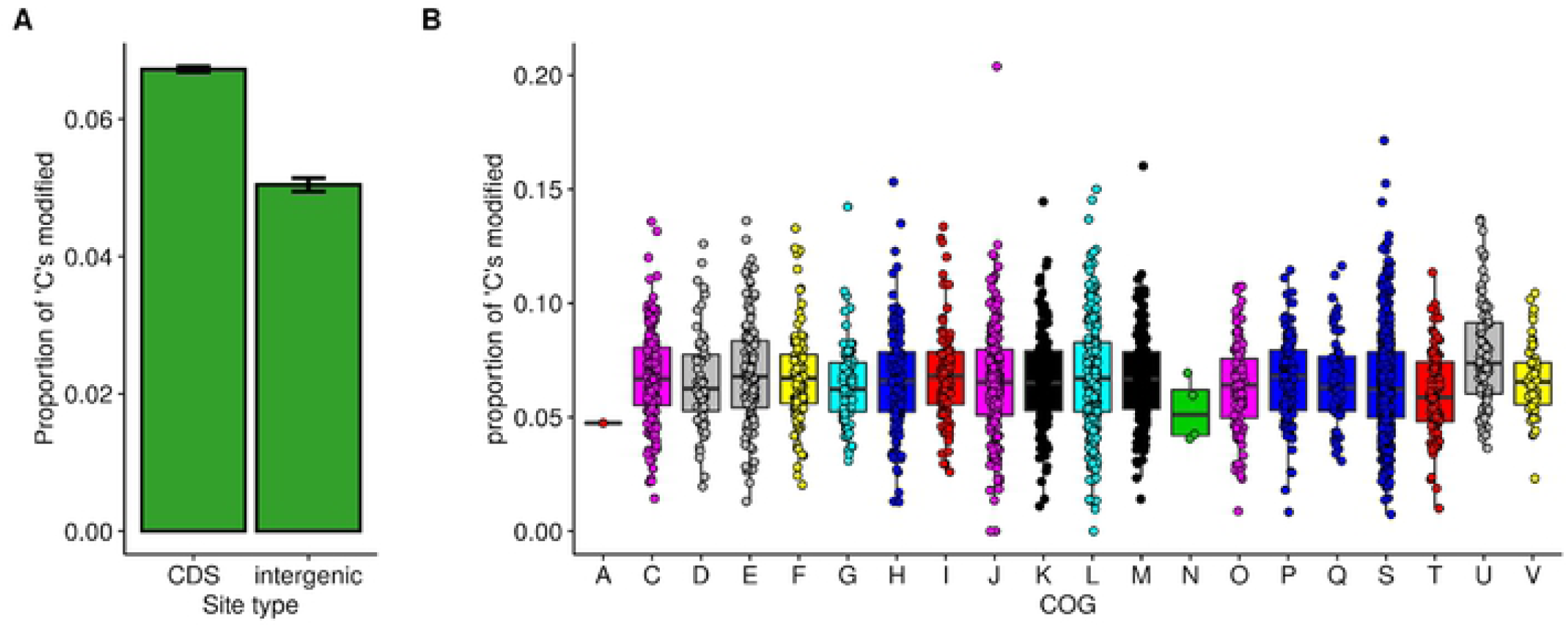
m4C methylation is not strongly associated with any functional class. (A) The proportion of ‘C’ nucleotides are with evidence for methylation in coding and intergenic sequences. Error bars represent a 95% confidence interval, calculated using the normal approximation of a binomial distribution. (B) Each point represents the proportion of ‘C’ nucleotides in a given gene that show evidence for methylation. The genes are grouped and shaded by the COG category (x-axis). The box plots summarise the distribution of this value across each COG category

In summary, this study has demonstrated that most variability in the *L. longbeachae* genome is from recombination and there have been large-scale rearrangements within the bacterial chromosome. Our 54 sg1 clinical isolates could be grouped into two highly related clades that persisted over time. The most genetically distinct clade consisted of isolates from only the Canterbury region but this could just reflect oversampling from this region and further sequencing of isolates from other regions is required. Most *L. longbeachae* isolates sequenced in this study were found to contain a plasmid. The plasmids showed high levels of recombination and horizontal gene transfer with evidence for both intra- and inter-species gene-flow. The genome of *L. longbeachae* was also highly modified, with m6A modifications being the most common and strongly associated with particular sequence motifs.

## Materials and methods

### Bacterial isolates, sequencing and genome assembly

A total of 60 isolates previously identified as *L. longbeachae* (including 57 serotyped as sg1 and 3 serotyped either as sg2 or undefined) were sequenced. Isolates were obtained from either the NZ *Legionella* Reference Laboratory (ESR, Porirua, New Zealand; n=39) or Canterbury Health Laboratories (CHL) culture collection (Christchurch, New Zealand; n=21). All isolates were derived from sporadic LD cases that occurred between 1993 and 2015 from 8 regions (S7 Fig) around the country and included the first NZ case in which *L. longbeachae* was successfully cultured from a patient specimen (LA93_171; S3 Table).

The isolates were grown on buffered-charcoal-yeast-extract (BCYE) agar at 37°C for 72 hours. DNA was extracted from each fresh culture using GenElute Bacterial Genomic kits (Sigma-Aldrich, MO, USA) according to the manufacturer’s instructions, including proteinase K and RNase treatments. Libraries were prepared from the genomic DNA using the Nextera XT kit (Illumina, San Diego, CA, USA) and were sequenced using Illumina MiSeq technology (2 × 250bp paired-end) and version 2 chemistry by New Zealand Genomics Ltd (NZGL; University of Otago, Dunedin, New Zealand). The quality of the raw reads was checked using FastQC (v. 0.11.4; https://www.bioinformatics.babraham.ac.uk/projects/fastqc/). They were mapped against PhiX using Bowtie2 (v. 2.0.2; [42]), and any that mapped to PhiX were removed from the SAM file, and read pairs were reconstructed using the SamToFastq.jar program from the Picard suite (v. 1.107; https://broadinstitute.github.io/picard/) using the default parameters. Any adaptors were removed through the “fastq-mcf” program (using the default parameters) from the ea-utils suite of tools (v. 1.1.2-621; https://expressionanalysis.github.io/ea-utils/). Finally the reads were quality trimmed using SolexaQA++ (v. 3.1.4; [43]) at a probability threshold of 0.01 and sorted on length to remove any sequences < 50 bp prior to assembly. Sequence reads from each isolate was assembled using the SPAdes (v. 1.2, [44]) *de novo* assembler in “careful” mode, with default settings.

### Sequenced strains analysed

Of the 60 isolates sequenced (S3 Table), 57 were found to be *L. longbeachae* sg1, two were sg2 and one had been mistyped and was found to be *Legionella sainthelensi*. Analyses were limited to the sg1 isolates but because poor sequence data were obtained for three genomes (2 from Auckland and 1 from Waikato), only 54 were included (Table 1). We also included the two other publically available complete genomes for *L. longbeachae* sg1, NSW150 (Australia; GenBank: NC_013861) and FDAARGOS_201 (USA; GenBank: NZ_CP020412) in our core genome and cluster of orthologous groups (COG) analyses.

**Table 1:** Number of isolates sequenced and analysed by year group and region

| Region | Year Group |  |  |  |  | Total |
| --- | --- | --- | --- | --- | --- | --- |
|  | 1993-1997 | 1998-2002 | 2003-2007 | 2008-2012 | 2013-2015 |  |
| Northland | - | 2 | - | - | - | 2 |
| Auckland | - | - | - | 2 | 9 | 11 |
| Waikato | 1 | - | 1 | - | - | 2 |
| Bay of Plenty | - | - | 1 | - | - | 1 |
| Rotorua | - | - | - | - | 1 | 1 |
| Manawatu | - | 1 | - | - | - | 1 |
| Canterbury | 6 | 1 | 9 | 14 | 4 | 34 |
| Otago | - | - | - | - | 2 | 2 |
| Total | 7 | 4 | 11 | 16 | 16 | 54 |

The reads of a further 65 previously published *L. longbeachae* isolates (Bioproject number PRJEB14754, SRA accession numbers ERS1345649 to ERS1345585; [17]) were downloaded and compared with our 54 sg1 isolates. However, 30 of these read sets were either of poor quality, aligning to less than 80% of our reference genome (F1157CHC; GenBank NZ_CP020894; [19]), or were not *L. longbeachae* sg1 isolates. These were excluded and the remaining 35 read sets were included in our global phylogenetic analyses (S6 Table).

### Complete New Zealand reference genome, gene prediction and annotation

To generate our own complete NZ reference genome, one isolate (F1157CHC; GenBank NZ_CP020894.2) was further sequenced using the PacBio RSII system (Pacific Biosciences, CA, USA) as previously described [19]. Briefly, the isolate was cultured (as above) and the DNA was extracted from fresh culture using the Blood and Cell Culture DNA Midi Kit (Qiagen, Hilden, Germany). One SMRTbell DNA library was constructed according to the manufacturer’s 20kb protocol, size selected using a 15 kb cut-off with BluePippin (Sage Science, MA, USA) and sequenced using P6-C4 chemistry and a 240 minute data collection time on one SMRT cell by the Doherty Institute for Infection & Immunity, University of Melbourne (Melbourne, Australia). A pre-assembly filter removed reads shorter than 500 bp or with a quality lower than 80%. The data was assembled using the HGAP2 assembly pipeline in SMRTanalysis v. 2.3.0. The minimum seed read length was 6000 bp and the Celera assembler stage assumed an approximate genome size of 5 Mb and allowed an overlap error rate of 0.06 and a minimum overlap length of 40 bp. The assembly was polished using Quiver and the MiSeq reads were mapped onto the final RSII assembly using Pilon (v. 1.20; [45]) to assess accuracy (99.99%). Gene prediction and annotation was performed using the NCBI Prokaryotic Genome Annotation Pipeline (2013).

### Genome architecture

#### Chromosome

In order to assess the genome architecture, F1157CHC was used as the basis for all analyses in which comparisons were made against a reference. The genome was visualized using Circos software (v. 0.69.3, [46]) and in this way, various other tracks of information described herein could be included on the plot. This included mapping the annotation prediction from PGAP, as well an overlay of the results of a functional annotation with the eggNOG web annotation server (see below), mapping of both the methylation results (see below) and recombinant regions detected with Gubbins (see below), SNP density of the comparative samples, and finally a visualization of the repeats within the F1157CHC genome using Reputer ([47, 48]).

The genome was analysed with the following Reputer parameters (number of best hits: 10000; minimum length: 30bp; and maximum Hamming distance: 3), and the output parsed through a MySQL database with a custom Perl script to generate the tracks to allow the links between all repeated regions to be visualized on the Circos plot. Of the four possible detectable repeats, only the forward (in blue) and palindromic (in red) repeats were detected, reverse and complement were not. Furthermore, depending on the Hamming distance between the two repeats, the links were coloured to show those with a smaller Hamming distance as a darker colour. In order to assess the overall genome architecture in comparison to other *L. longbeachae* genomes, the MAUVE plugin within Geneious (v. 9.1.8) was used to visualize the F1157CHC genome against NSW150 and FDAARGOS_201.

#### Plasmid

The five *L. longbeachae* reference plasmids pB1445CHC_73k (CP045304), pB1445CHC_150k (CP045305), pB41211CHC_76k (CP045307), pLLO-F1157CHC (CP020895) and pNSW150 (NC_014544) were aligned and visualised using MAUVE plugin within Geneious (v.9.1.8).

### Genetic diversity of *L. longbeachae* sg1 clinical isolates

#### Core genome and COG analyses

The eggNOG-mapper [28, 49]) webserver (http://eggnog-mapper.embl.de/) was used to annotate the F1157CHC PGAP-derived amino acid sequences. Default parameters were used for the annotation. The Prokka pipeline (v. 1.12; [50]) was used to annotate our draft isolates using default parameters. The Prokka-generated GFF files were analysed with Roary using default parameters, and the comparison script roary_plots.py was used to visualize the output. FastGeP was used with default parameters to perform a whole genome MLST analysis of the 56 isolates, which meant that the generated allele sequences were searched with BLAST+ at an identity threshold ≥ 80%. CP020894 was used as the reference genome for this analysis. SplitsTree (v.4.15.2, [51, 52]) was used to convert the FastGeP Nexus file into a Newick file (as a Neighbour-joining tree) for visualization and annotation in iTOL with the inclusion of metadata for region and the sample type.

#### Single nucleotide polymorphism identification

Single nucleotide polymorphisms (SNPs) were identified using Snippy v2.6 (https://github.com/tseeman/snippy). Snippy is a pipeline that uses the Burrows-Wheelers Aligner [53] and SAMtools [54] to align reads from different isolates to our reference genome (F1157CHC; GenBank CP020894) and FreeBayes [55] to identify variants among the alignments. Gubbins was used to remove areas of recombination on the full alignment [56]. Such data was visualised on the Circos plot of F1157CHC described above.

#### Antimicrobial Resistance and Virulence Genes

The contigs of each isolate were screened and acquired resistance and virulence genes were identified using ABRIcate (v. 2; https://github.com/tseemann/abricate).

### Determination of the geo-temporal and phylogenetic relationships

#### Ancestral state reconstruction and phylogenetic analysis

Snippy v2.6 (https://github.com/tseeman/snippy/) was used to align reads of 54 New Zealand *L. longbeachae* isolate to the reference genome *Legionella longbeachae* F1157CHC (CP020895) and identify SNPs. Gubbins was used to remove areas of recombination [56]. SNPs were exported into BEAUti v2.5 to create an Extensive Markup Language (xml) file for BEAST v2.5 [57]. The *Legionella longbeachae* reference genome consists of 1,306,681 adenine, 765,717 cytosine, 772,189 guanine and 1,298,289 nucleotides. These nucleotides were added as constant sites to keep the model representative of *L. longbeachae*. bModelTest [58] was used to choose the substitution model. Multiple molecular clock and tree models were trialed. Nested sampling (NS) [59] was used to select the model (S3 File) [60]. A 121123 Generalised Time-Reversible (GTR) [61] model was used to model nucleotide substitutions, an Extended Bayesian Skyline [62] was used to model the effective population size, and an uncorrelated relaxed clock [63] was used to model the molecular clock and was calibrated by tip dates. The .xml file was run in BEAST for 100 million steps, three times with different starting seeds, before LogCombiner was used to combine the runs with a 10% burn-in. Tracer v1.6 (Rambaut et al, http://beast.bio.ed.ac.uk/Tracer) was used to visualise the results. TreeAnnotator v2.6 was used to form a maximum clade credibility tree from the trees produced using BEAST. Evolview v2 [64] was used to visualise and edit the tree. The raw reads of the 54 New Zealand *L. longbeachae* isolates were uploaded to NCBI (PRJNA562040), along with the four excluded from analysis.

#### Plasmid Analyses

The assemble contig reads of the 54 New Zealand *L. longbeachae* sg 1 isolates were aligned to the five *L. longbeachae* reference plasmids pB1445CHC_73k, pB1445CHC_150k, pB41211CHC_76k, pLLO-F1157CHC and pLLO-NSW150, using Burrows-Wheeler aligner [53]. For each read set, the proportion of the plasmid with a read depth of ten or higher was calculated using samtools v1.9 [54]. The plasmid sequences were annotated using Prokka v1.14.4 [50] and a dendrogram of gene presence-absence was formed using Roary v3.11.2 [29], before the sequences were aligned using EasyFig v2.2.4 [65].

#### Global *L. longbeachae* isolates

The reads of 64 previously published *L. longbeachae* isolates [17] were downloaded and were compared with the 54 New Zealand isolates using the SNP-identification method described above. Of these read sets, 29 were of poor-quality, aligning to less than 80% of the reference, or were distantly related from the rest of the isolates, indicating that they were not *L. longbeachae* sg1. These isolates were excluded from analysis. In total, 89 *L. longbeachae* isolates from New Zealand and United Kingdom were investigated. RaxML [66] was used to form a maximum likelihood tree of the isolates based on their SNP data and was visualised using EvolView v2.

### *L. longbeachae* Methylome

Methylated bases were detected for isolate F1157CHC, using the “RS_Modification_and Motif Analysis” protocol implemented in SMRTAnalysis v2.3.0 using the SMRTbell DNA library described above as input. This pipeline takes advantage of BLASR (v1, [67]) to map sequencing reads to the assembled genome and MotifFinder v1 to identify sequence motifs associated with particular modifications. The resulting files were submitted to REBASE [68] along with our annotated reference genome to identify protein coding genes that may be responsible for the inferred methylation patterns.

The distribution of methylated bases on the reference genome, and with regard to genomic features was analysed using bedtools (v2.25.0, [69]) and the R statistical language (v3.4). We tested for differences in methylation rate between genes of different functional classes using anova, as implemented in R. A complete record of the code used to perform statistical analyses and visualisation of the methylome data is provided in (S4 File).

## Acknowledgements

This work was funded through a University of Otago Research Grant and the PacBio sequencing of our complete reference genome was funded by a Millenium Science SMRT Grant. The PacBio sequencing service was provided by the Doherty Institute for Infection and Immunity, University of Melbourne, Australia. The Illumina MiSeq sequencing service was provided by the Massey Genome Service (Massey University, Palmerston North, New Zealand), as part of NZ Genomics Ltd, Dunedin, New Zealand.

We thank David Harte from the Legionella Reference Laboratory, ESR, Porirua, New Zealand and Canterbury Health Laboratories, Christchurch, New Zealand for supplying the clinical *L. longbeachae* isolates.

## Supporting Information

Supplementary Figure 1: Colours of functional COG categories as depicted in the Circos plot-figure 2.

Supplementary Figure 2: Mauve alignment of *Legionella* plasmids.

Supplementary Figure 3: NeighbourNet tree of 54 *L. longbeachae* clinical isolates based on 1,271 core SNPs.

Supplementary Figure 4: Dendrogram of *L. longbeachae* reference plasmid sequences based on gene presence-absence, and alignment of sequences.

Supplementary Figure 5: Maximum likelihood tree of 89 *L. longbeachae* isolates from New Zealand and the United Kingdom.

Supplementary Figure 6: Regional Origin of the Sequenced Isolates. The number of isolates sequenced from each region is shown in brackets.

Supplementary Table 1: Mauve collinear blocks Supplementary Table 2: eggNOG annotations/output

Supplementary Table 3: Meta- and genome summary data from Prokka of the *Legionella longbeachae* clinical isolates sequenced and analysed in the current study.

Supplementary Table 4: MIC_90_ (mg/L) values for *L. longbeachae* isolates by broth dilution (BYE; Isenman et al., 2018).

Supplementary Table 5: Virulence factor genes identified the L. longbeachae isolates sequenced in the current study.

Supplementary Table 6: Sg1 clinical and environmental isolates from Bacigalupae *et al*., 2017^†^ that were included in the global phylogenetic analysis.

Supplementary File 1: R-Plots-Roary outputs file

Supplementary File 2: FastGeP matrix in iTOL output file

S3 File: Molecular Clock and Tree Model Trialing

S4 File: Code used to perform statistical analyses and visualisation of the methylome data.

